# G protein-coupled estrogen receptor is not required for sex determination or ovary function in zebrafish

**DOI:** 10.1101/373092

**Authors:** Camerron M. Crowder, Shannon N. Romano, Daniel A. Gorelick

**Affiliations:** Department of Pharmacology & Toxicology, University of Alabama at Birmingham, Birmingham Alabama, USA; Center for Precision Environmental Health, Department of Molecular & Cellular Biology, Baylor College of Medicine, Houston, Texas, USA

**Author notes:** present address: Department of Cell, Developmental and Regenerative Biology, Icahn School of Medicine at Mount Sinai, New York, New York, USA. present address: Department of Cell, Developmental and Integrative Biology, University of Alabama at Birmingham, Birmingham Alabama, USA.

## Abstract

Estrogens regulate vertebrate development and function through binding to nuclear estrogen receptors alpha and beta (ERα, ERβ) and the G protein-coupled estrogen receptor (GPER). Studies in mutant animal models demonstrated that ERα and ERβ are required for normal ovary development and function. However, the degree to which GPER signaling contributes to ovary development and function is less well understood. Previous studies using cultured fish oocytes found that estradiol inhibits oocyte maturation in a GPER-dependent manner, but whether GPER regulates oocyte maturation *in vivo* is not known. To test the hypothesis that GPER regulates oocyte maturation *in vivo*, we assayed ovary development and function in *gper* mutant zebrafish. We found that homozygous mutant *gper* embryos developed into male and female adults with normal sex ratios and fertility. Adult mutant fish exhibited normal secondary sex characteristics and fertility. Additionally, mutant ovaries were histologically normal. We observed no differences in the number of immature versus mature oocytes in mutant versus wild-type ovaries from both young and aged adults. Furthermore, expression of genes associated with sex determination and ovary function were normal in *gper* mutant ovaries compared to wild type. Our findings suggest that GPER is not required for sex determination, ovary development or fertility in zebrafish.

## INTRODUCTION

Estrogens influence a wide range of physiological processes in reproductive and non-reproductive tissues in zebrafish [1-6]. Estrogens act by binding to nuclear estrogen receptors (ERα, ERβ), ligand dependent transcription factors that directly regulate gene expression [7-9]. Estrogens also activate the G protein-coupled estrogen receptor (GPER/GPR30) to elicit downstream signaling cascades [10-12]. In the zebrafish ovary, two ERα paralogue genes (*esr2a* and *esr2b*) are essential in reproduction, folliculogenesis and maintenance of ovarian tissue, with mutations in both receptors resulting in female to male sex reversal [1]. However, less is understood regarding the role of GPER in sex determination, ovarian development and oogenesis in vertebrates.

Sexual determination in zebrafish is unique in that laboratory strains lack a sex chromosome and no sex-determining gene has been identified. Elevated temperatures and stress are associated with masculinization, but these results are conflicting [13-15]. It appears that multiple sex-related genes interact as a network to establish sex, a concept known as polygenic sex determination [16-18]. Several genes promote ovary differentiation and development, including aromatase (*cyp19a1a*) and forkhead box L2 (*foxl2a*, *foxl2b*) [17, 19-22]. Conversely, expression of anti-Mullerian hormone gene (*amh*) promotes testes differentiation and spermatogenesis [21, 23-25]. Additionally, exposure of juvenile zebrafish to exogenous estrogens can bias sex differentiation in favor of fully functioning and fertile females [23, 26]. Although levels of estrogens are important for female sex determination and differentiation, the role of GPER in zebrafish sex determination and differentiation is unknown.

GPER was shown to regulate oocyte maturation *in vitro*. In carp (*Cyprinus carpio*) and zebrafish, maturation of cultured oocytes is regulated by estrogens and progestogens, where pharmacologic and RNA interference approaches demonstrated that estrogens inhibit maturation via GPER while progestogens promote maturation via membrane progesterone receptors [5, 27-30]. In ovaries cultured from golden hamsters (*Mesocricetus auratus*), estradiol promoted the formation of primordial follicles in a GPER-dependent manner [31].

While evidence from *in vitro* studies demonstrates that GPER regulates oocyte maturation and ovarian follicle development, evidence from *in vivo* studies suggests that GPER is not required for ovary development. *Gper* mutant mice showed no defects in fertility or gonad development [32, 33].

In carp and zebrafish ovaries, *gper* mRNA was detected in pre-vitellogenic, late-vitellogenic and post-vitellogenic oocytes but was not detected in ovarian follicle cells [27, 34]. Thus, *gper* expression in teleost ovaries is consistent with a cell-autonomous role in oocyte maturation. To better understand the role of GPER in the development and function of ovaries, we examined ovary histology, fertility and gene expression in adult wild-type and *gper* mutant zebrafish. Our findings suggest that GPER does not influence sex determination, gonad development or fertility in zebrafish.

## METHODS

### Animal care and husbandry

Zebrafish were housed at the University of Alabama at Birmingham (UAB) zebrafish facility in a recirculating water system (Aquaneering, Inc., San Diego CA) and kept at 28.5°C on a 14-h light, 10-h dark cycle. Zebrafish embryos were raised in E3B (60x E3B: 17.2 g NaCl, 0.75 g KCl, 2.9g CaCl_2_-H_2_0, 2.39 g MgSO_4_ dissolved in 1 L Milli-Q water; diluted to 1x in 9 L Milli-Q water plus 100 μl 0.02% methylene blue) and housed in an incubator at 28.5°C on a 14-h light, 10-h dark cycle, until 5 dpf. Wild-type zebrafish were AB strain [35]. *gper^uab102^* mutant zebrafish, containing a 130 basepair deletion in the open reading frame, were used as described [2]. All procedures were approved by the UAB and Baylor College of Medicine Institutional Animal Care and Use Committees.

### Genomic DNA isolation and genotyping analysis

Fin biopsies from wild type and *gper* mutant adults were digested in 100 μL lysis buffer (10 mM Tris pH 8.3, 50 nM KCl, 0.3% Tween 20) with 1 μL proteinase K (800 U/mL, New England Biolabs) per well in 96-well plates. Samples were incubated at 55°C for 8 hours to extract genomic DNA, followed by 10 minute incubation at 98°C to inactivate proteinase K. Samples were stored at −20°C. Genotyping was performed using polymerase chain reaction and gel electrophoresis as described [2].

### Secondary sexual characteristics and sex ratios

Heterozygous *gper^uab102/+^* adults were bred to each other and offspring were raised to adulthood. At approximately 4-months of age progeny were genotyped and secondary sexual characteristics were examined visually. Zebrafish sex was determined by examining multiple secondary sexual characteristics including body shape, anal fin coloration, and presence or absence of a genital papilla (Figure 1, 2,)[36]. Mendelian ratios were determined via genotyping. The total number of males and females for each genotype were counted and compared to heterozygous and wild-type animals from the same clutch (Table 1, Figure 1).

**Figure 1.**
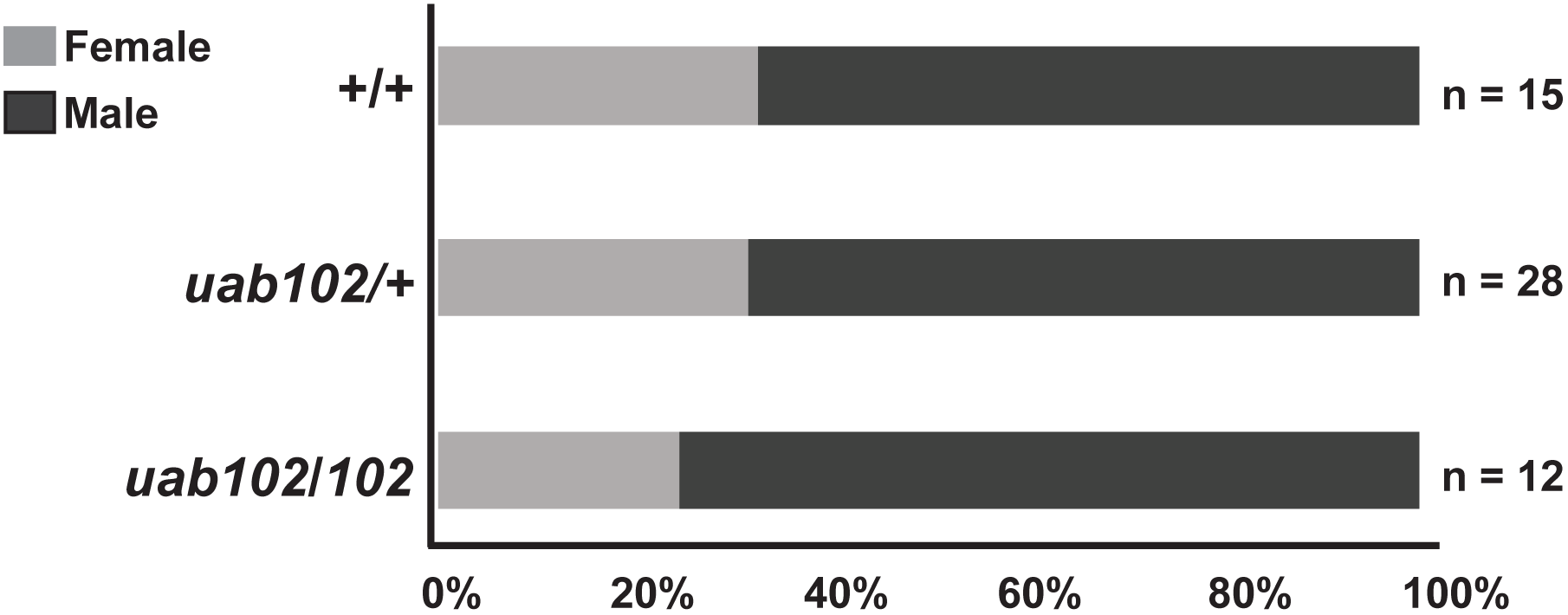
*gper^uab102/102^* mutant adults display normal sex ratios. Percentage of male and female progeny from crosses between *gper^102/+^* males and females. Offspring were observed to have 1:2:1 Mendelian ratios and sex ratios were similar across all three genotypes. Number of fish per genotype is included to the right of the graph.

**Figure 2.**
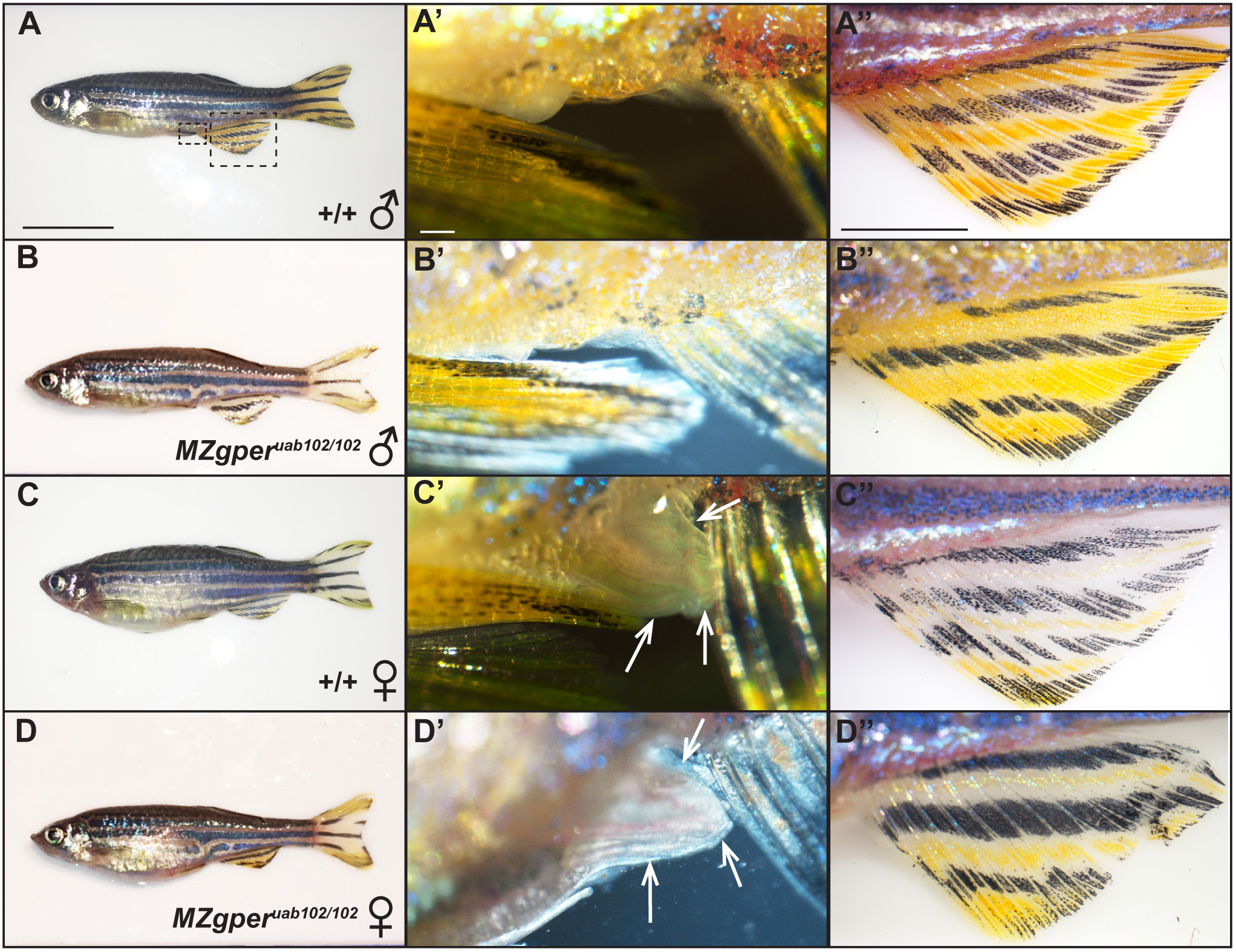
*MZgper^uab102/102^* mutant males and females display normal secondary sex characteristics. Male wild-type fish have a slender body shape (A), lack a genital papilla (A’) and have a yellow anal fin (A’’). Similarly, *MZgper^uab102/102^* mutants have a slender body shape (B), no genital papilla (B’) and a yellow anal fin (B’’). Wild-type females (C) and *MZgper^uab102/102^* females (D) have a rounded body shape with prominent protruding abdomen, large extended genital papilla (arrows in C’, D’) and whiter anal fin (C’’, D’’). (A’, A’’) High magnification images of boxed areas in A, which corresponds to all other similar images. Scale bars: 1 cm (A-D), 100 μm (A’-D’), 1000 μm (A’’-D’’). 10 females and 8 males per genotype were examined.

**Table 1.** Mendelian and sex ratios in progeny derived from *gper^uab102/+^* crosses. A single population of offspring derived from harem breeding among male and female *uab105/+* heterozygotes. Total number of individual fish within each genotype is indicated, followed by percent of fish with indicated genotype. Numbers of male and females for each genotype are included in brackets, with sex based on presence of testis (♂) or ovary (♀).

|  | **+/+** | ***uab102/+*** | ***uab102/102*** |
| --- | --- | --- | --- |
| **Population 1** | 15 (27%)<br>[10 ♂, 5 ♀] | 28 (51%)<br>[19 ♂, 9 ♀] | 12 (22%)<br>[9 ♂, 3 ♀] |

### Histology

All histology experiments were performed on wild type or maternal zygotic homozygous *gper* mutant adult fish (MZ*gper^uab102/102^*). *MZgper^uab102/102^* individuals were generated by crossing homozygous (*gper^uab102/102^*) females to homozygous males. Whole-body 4-month and 10-month old adult *MZgper^uab102/102^* mutant and wild-type females (n = 4 females per genotype) were selected for histology based on secondary sexual characteristics and breeding trials. Fish were anesthetized in iced 0.2 mg/mL tricaine, decapitated with a razor blade, followed by a 2-day fixation in 4% formaldehyde. Specimens were either processed, embedded and sectioned as described [36] or shipped on ice to HistoWiz Inc. (New York, NY) where they were processed, embedded, sectioned (5 microns) and stained with hematoxylin and eosin (H & E) as follows: Processing of tissues into paraffin blocks was accomplished using an automated Peloris II tissue processor (Leica Biosystems, Buffalo Grove, IL) and embedded in Histoplast PE (Thermo Scientific). Tissues were dehydrated using the following protocol: 50% ethanol (15 minutes, 45°C), 70% ethanol (15 minutes, 45°C) × 2, 90% ethanol (15 minutes, 45°C), 90% ethanol (15 minutes, 45°C), 90% ethanol (30 minutes, 45°C), 100% ethanol (45 minutes, 45°C), 100% xylene (45 minutes, 45°C), Parablock wax (30 minutes, 65°C) × 2 (Leica Biosystems) and Parablock wax (45 minutes, 65°C). Embedded specimens were cut into 5-micron sections and adhered to charged slides in a water bath. Slides were heated to 65°C for 10 minutes in an oven to melt paraffin and fully-adhere sections. Slides were rehydrated and stained using the a Tissue-Tek Prisma (Sakura) or by hand in glass containers using the following protocol: 100% xylene (5 minutes) × 2, 100% ethanol (2 minutes) × 2, 95% ethanol (1 minute), deionized (DI) H_2_O (1 minute), hematoxylin (1 minute), DI H_2_O (1 minute), defining solution (45 seconds) (Define MX-aq; Leica Biosystems), DI H_2_O (1 minute), bluing agent (Fisher Scientific), DI H_2_O (1 minute), 95% ethanol (15 seconds), eosin (30 seconds), 95% ethanol (2 minutes), 100% ethanol (3 minutes) × 2 and 100% xylene (5 minutes) × 2. Brightfield images of sections were obtained using a Zeiss Axio Observer.Z1 microscope with a Zeiss Axio MRc5 camera with 10x and 20x objectives (Figures 4). In addition, specimens processed at HistoWiz were imaged using a Leica Aperio AT2 slide scanner (Figure 5). A one-way ANOVA with Tukey’s test for multiple comparisons was performed to determine significance (*p* ≤ 0.05) in total number of oocyte per stage, all statistics were completed using GraphPad Prism software (version 7.0).

**Figure 3.**
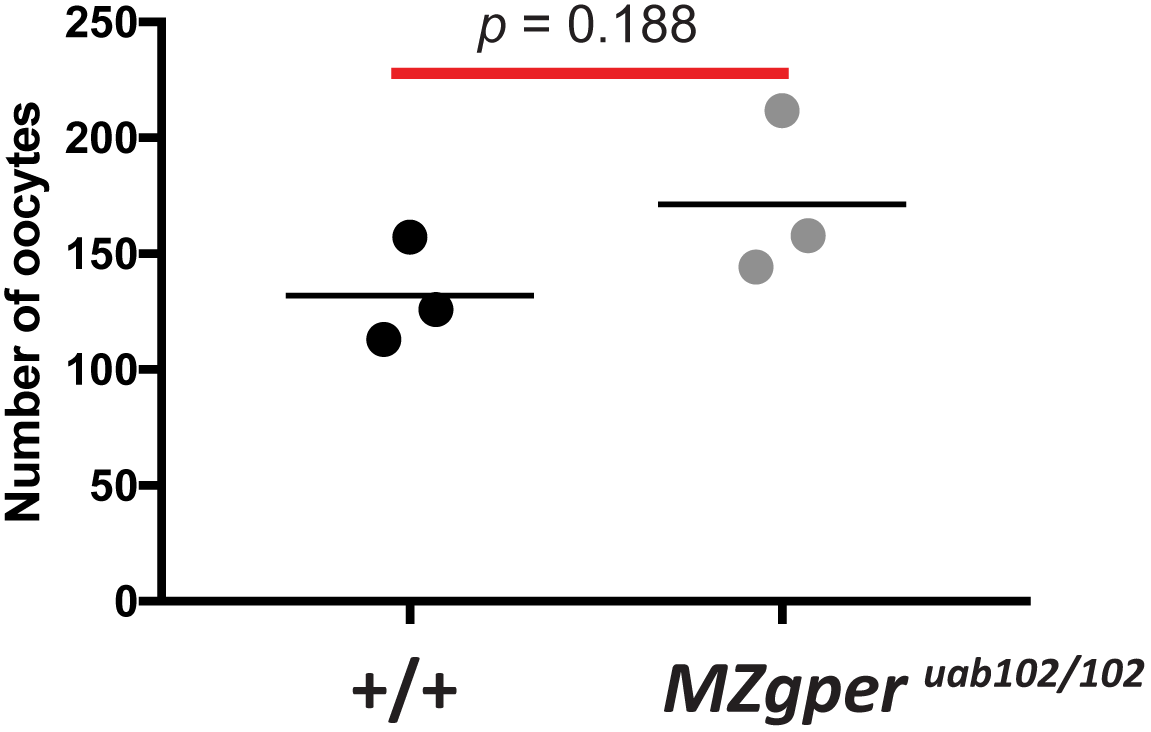
*MZgper^uab102/102^* mutants release on average a similar numbers of oocytes to wild-type fish during natural breeding trials. *MZgper* and wild-type 10-month old adult females were mated with wild-type males in 1:1 ratio (n = 10 fish per trial). Each dot represents the average number of oocytes released by spawning females in each mating trial (n =3 trials). *P* = 0.188, unpaired t-test.

**Figure 4.**
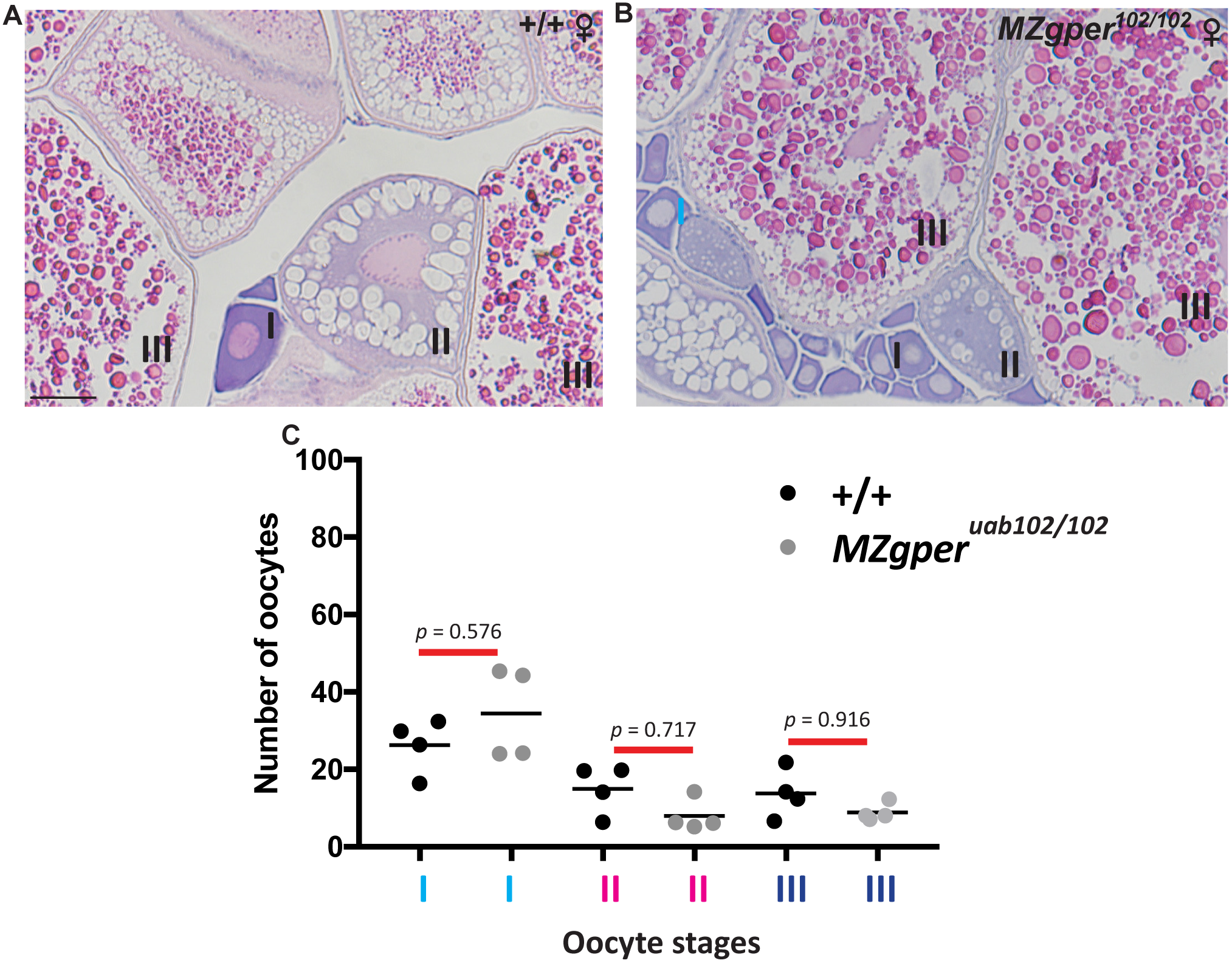
4-month old *MZgper^uab102/102^* and wild-type ovaries contain similar quantities of all stages of oocytes. H & E staining of adult 4-month old wild-type (A) and *MZgper* (B) ovaries. Number of oocytes of each stage (I, II & III) are shown in (A, B) and quantified in (C) with *p*-values from one-way ANOVA with Tukey’s test for multiple comparisons. Each dot represents average number of oocytes per histological section per fish (n = 4 fish per genotype, 15 sections per fish). Scale bar = 200 μm

**Figure 5.**
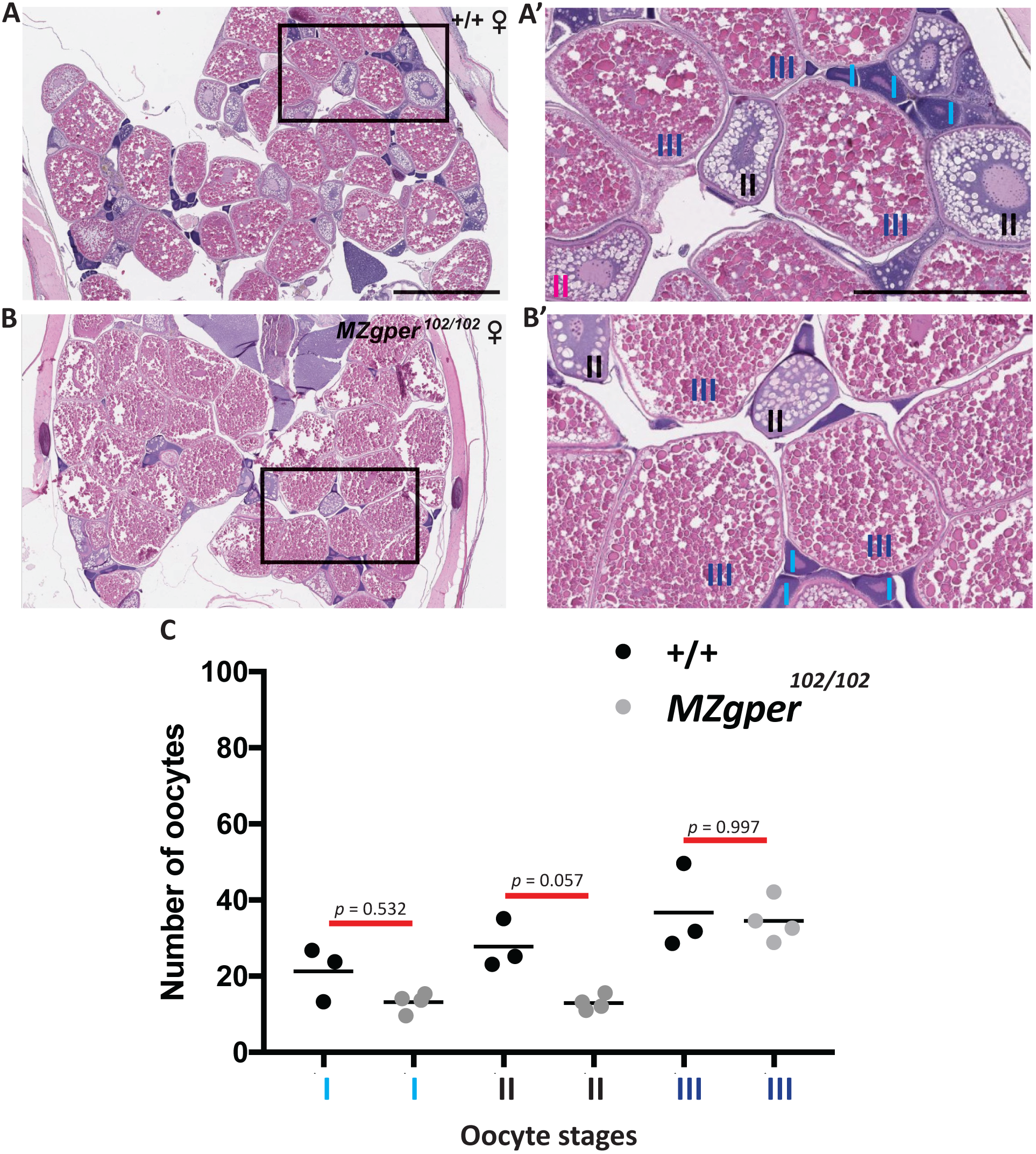
10-month old *MZgper^uab102/102^* and wild-type ovaries contain similar quantities of all stages of oocytes. H & E staining of adult 10-month old wild type and *MZgper^uab102/102^* ovaries (A, B). Boxed areas are shown at higher magnification (A’, B’). Number of oocytes according to stage (I, II & III) are labeled (A’, B’) and quantified (C) with p-values from one-way ANOVA with Tukey’s test for multiple comparisons. Each dot represents average number of oocytes per histological section per fish (n = 3-4 fish per genotype, 15 sections per fish). Scale bars = 1 mm (A, B), 500 μm (A’, B’).

### Breeding trials

*MZgper^uab102/102^* mutant females were bred once every two weeks with wild-type males (1 female: 1 male) and compared to wild-type crosses (1 female: 1 male) (n = 10 adult females per genotype). The number of individuals that spawned and the total number of oocytes released were counted after each breeding trial (Table 2, Figure 3). An unpaired t test was performed to determine significance, defined as *p* ≤ 0.05, in number of oocytes released. Statistical analyses were performed using GraphPad Prism software (version 7.0).

**Table 2.** *MZgper^uab102/102^* spawn as frequently and release similar numbers of oocytes as wild-type females. 10-month old adult *MZgper^uab102/102^* or wild-type females were crossed 1:1 with wild-type males (n =10 individuals per genotype). The percent of pairs that spawned was determined by counting the number of females that released oocytes in each trial. For each trial of 10 crosses, the average number of oocytes released is shown (mean ± standard deviation).

| Genotype of female paired with wild-type male | Trial 1 | Trial 2 | Trial 3 |
| --- | --- | --- | --- |
| **Wild type** | 60% spawned<br>126 oocytes ± 49 | 60% spawned<br>158 oocytes ± 55 | 70% spawned<br>113 oocytes ± 88 |
| ***MZgper*<sup>uab102/102</sup>** | 50% spawned<br>157 oocytes ± 65 | 50% spawned<br>212 oocytes ± 61 | 70% spawned<br>144 oocytes ± 50 |

### RNA extraction and quantitative PCR

Total RNA was extracted from four dissected ovaries of 10-month old adult *MZgper^uab102/102^* and wild-type zebrafish according to manufacturer’s guidelines for the TRIzol RNA isolation kit (Life Technologies, Carlbad, CA). Ovaries were homogenized individually in 500 μL TRIzol and DNase-treated with a TURBO DNA-free kit (Ambion) to remove genomic contamination. Quality of RNA was checked on a 1% agarose gel and quantity of RNA was determined using a Nanodrop ND-1000 ultraviolet-Vis spectrophotometer (Thermo Scientific). Expression of genes previously shown to be involved in sex differentiation were examined with quantitative reverse transcription PCR (qRT-PCR): *cyp19a1a*, *foxl2a, amh* and *rp113a* as a reference gene (Supplemental Table 1) [17, 37, 38]. Complementary DNA (cDNA) was synthesized via reverse transcription according to manufacturer’s guidelines for the RETROscript First Strand Synthesis Kit (Fisher Scientific). Quantitative RT-PCR reactions were completed using SsoAdvanced Universal SYBR Green Supermix Kit (Bio-Rad Laboratories, Inc.) and run on a Bio-Rad CFX96 Touch Real-Time PCR Detection System using the following protocol: 98°C for 1 minute, followed by 34 cycles of 98°C for 10 seconds, 60°C or 61°C for 20 seconds, 72°C for 45 seconds and then 72°C for 2 minutes. Reactions were run in technical quadruplicates on 96-well plates and transcript quantification was measured using CFX Manager version 3.1 software (Bio-Rad Laboratories, Inc.). Cq expression values were averaged across technical replicates, primer efficiencies were measured and expression fold change values were calculated according to the ∆∆Ct method [39]. An unpaired t test was performed on relative expression levels (2^−∆Ct^) for individual biological replicates (n = 3-4 per gene per genotype) to determine significant differences in gene expression between mutants and wild type. For the graph in Figure 6, relative expression was normalized to wild type mean for each gene.

**Figure 6.**
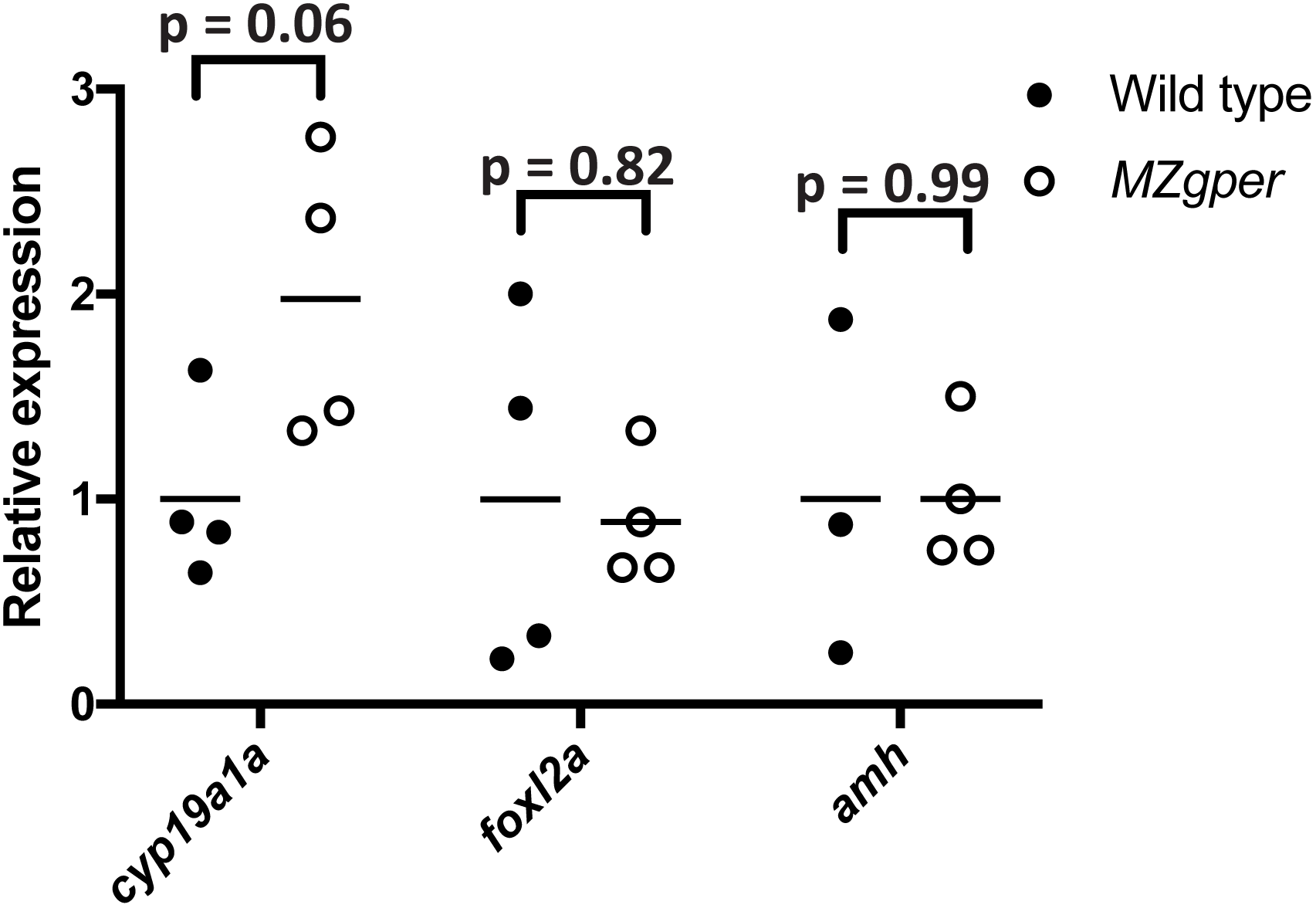
Quantitative RT-PCR of genes associated with sex determination and development in *MZgper^uab102/102^* mutant and wild-type ovaries. **(A)** qPCR analysis of *cyp19a1a*, *foxl2a* and *amh* mRNA in wild-type and *MZgper* mutant ovaries. Circles represent independent biological replicates, horizontal line is the mean, wild type mean set to 1 for each gene. Unpaired t-test showed no significant difference between gene expression in wild type and mutants.

## RESULTS

### Male and female ^*gperuab102/102*^ mutants display normal sex ratios compared to wild-type and heterozygous siblings

To determine if GPER is involved in sex determination in zebrafish we crossed *gper^uab102/+^* heterozygous males and females and compared the number of male and female offspring across genotypes. Males were the dominant sex in all genotypes (+/+, *gper^uab102/+^, gper^uab102/102^*), which is consistent with previous studies demonstrating that approximately 70% of wild type fish are male in the UAB zebrafish research facility [40]. There was a 67% male bias in wild-type fish, a 68% male bias in heterozygous fish and a 75% male bias in homozygous *gper^uab102/102^* fish (Table 1, Figure 1). Therefore, we conclude that GPER does not regulate sex determination in zebrafish.

### *gper* mutant males and females display normal secondary sexual characteristics compared to wild-type fish

Following our sex-ratio studies, we attempted to grossly assay fertility and ovary function by naturally breeding homozygous mutant females with homozygous mutant males. We found that homozygous mutant *gper* fish of both sexes are fertile. Therefore, we maintained *gper* mutants as homozygotes in our zebrafish colony by crossing homozygous adults to each other and raising maternal-zygotic homozygous embryos to adulthood (*MZgper^uab102/102^*). To evaluate if *gper* is involved in the physical appearance of males and females we examined secondary-sexual characteristics in *MZgper^uab102/102^* mutants compared to wild type. Wild-type males have a slender body shape, yellow anal fin and no genital papilla [36, 41]. Similarly, *MZgper^uab102/102^* mutant males displayed these same physical attributes (Figure 2A, B). In contrast, wild-type females display a rounded abdomen, a lighter colored anal fin and a large protruding genital papilla [36, 41]. Similarly, *MZgper^uab10/102^* mutant females displayed these same physical characteristics (Figure 2 C, D). Therefore, we conclude that *gper* does not influence the development of secondary sex characteristics.

### *gper* mutant females spawn as frequently and release a similar number of oocytes on average as wild-type adult females

To examine if *gper* is involved in regulating fertility, we measured spawning frequency and the total number of oocytes released by *MZgper^uab102/102^* and wild-type females during three mating trials. On average, 63% of female wild-type and 56% of female *MZgper^uab102/102^* spawned during mating trials (10 fish per genotype per trial, n = 3 trials, Table 2, Figure 3). *MZgper^uab102/102^* females released an equivalent number of oocytes as did wild-type females. The average number of oocytes released by wild-type females was 131 ± 32 (SD), compared to 171 ± 35 released by *MZgper^uab102/102^* mutant females (Table 2, Figure 3, *p* = 0.188, unpaired t test). Therefore, we conclude that *gper* does not influence spawning or oocyte release.

### *gper* mutant female ovaries are structurally similar to wild type at 4-months and 10-months of age

To evaluate if *gper* mutant ovaries show a defect in ovary development or oocyte maturation, we examined ovaries from 4-month old adult ovaries fish and counted the number of stage I, II and III oocytes present. *MZgper^uab102/102^* mutant ovaries were histologically similar to wild-type ovaries, with no significant difference in the number of stage I, II or III oocytes in mutants versus wild type (Figure 4). Ovary function declines with age [42], and it is possible that *gper* could influence the maintenance of normal ovary function, a phenotype that would not be apparent in young, 4-month old adults. To test whether *gper* influences ovary function in aged animals, we compared ovaries from mutant and wild type 10-month old zebrafish. We observed no difference in gross ovary organization and H & E staining between wild type and mutants. We also observed no significant differences in the number of stage I, II or III oocytes between 10-month old wild type and mutant zebrafish (Figure 5). Therefore, we conclude that *gper* does not regulate oocyte maturation or ovary function in laboratory zebrafish.

### Genes associated with sex determination and gonad function are not differentially expressed between *gper* mutant and wild-type ovaries

It is possible that GPER influences sex determination and ovary function by regulating gene expression. For example, *androgen receptor* mutant zebrafish have grossly histologically normal ovaries in adulthood, but display abnormal expression of sexually dimorphic genes associated with gonad function [36]. To determine whether GPER is involved in regulation of sexual determination or ovary differentiation and function we compared levels of three genes associated with sexual differentiation and maintenance: *cyp19a1a* and *foxl2a*, which promote and maintain ovary differentiation and function, and *amh*, which promotes testes differentiation [17, 24]. Expression levels of all three genes were not significantly different in *gper* mutants and wild-type adult zebrafish ovaries (Figure 6; *cyp19a1a* 2.01 ± 0.49 *MZgper* vs wild type, *foxl2a* 1.22 ± 1.53, *amh* 1.32 ± 1.50; p>0.05 in all cases, t test). Therefore, we conclude that GPER is not required for normal expression of genes associated with sex determination and maintenance.

## DISCUSSION

Our results suggest that GPER is not necessary for ovarian development and function in zebrafish. *gper* mutant ovaries appeared normal and contained all stages of oocytes in similar quantities to wild-type fish. Fertility was not affected in *gper* mutant females, which released a similar number of oocytes during natural mating trials and bred at a similar frequency to wild-type females. We also found that GPER does not influence sex determination or secondary sexual characteristics. *gper^uab102/102^* mutant embryos developed into male and female adults in similar ratios as wild-type and heterozygous clutch mates. Adult *gper^uab102/102^* males and females displayed normal sexually dimorphic body shape and anal fin coloration. *gper^uab102/102^* females possessed a large protruding genital papilla, similar to wild-type females. Expression of three genes important for sex determination or ovary maintenance (*cyp19a1a*, *foxl2a*, *amh*) were expressed at similar levels in *gper* mutant and wild-type ovaries, suggesting that these genes are not regulated directly or indirectly by GPER.

Our results are consistent with results from *Gper* mutant mice [32, 33]. However, our results contradict previously published *in vitro* results. In several fish species including zebrafish, estrogens were shown to inhibit meiotic maturation of cultured oocytes in a GPER-dependent manner [5, 27, 43, 44], yet we did not observe an oocyte maturation defect in *gper* mutant zebrafish. There are several explanations that could account for this discrepancy. One possibility is that assaying oocyte maturation *in vitro* does not reflect oocyte maturation *in vivo* with high fidelity. Studies examining the effects of estrogens on oocyte maturation *in vitro* use oocytes separated from follicle cells. In contrast, in the ovary, oocytes interact with follicle cells, extracellular matrix and circulating signaling molecules that could contribute to maintenance of meiotic arrest and make GPER unnecessary for maintenance of meiotic arrest in oocytes *in vivo*.

Another possibility is that cell signaling pathways compensate for lack of GPER signaling *in vivo*. Evidence suggests that, *in vitro*, GPER promotes meiotic arrest by increasing cGMP and cAMP levels [5, 45]. In the absence of GPER *in vivo*, perhaps another GPCR can compensate for GPER and drive an increase in cGMP and cAMP levels. Alternatively, there could exist novel or previously unappreciated estrogen receptors that contribute to estradiol-mediated inhibition of oocyte maturation in the absence of GPER.

A third possibility is that the *gper^uab102^* mutation is a hypomorph or is rescued by genetic compensation. The *uab102* allele lacks 130 basepairs from the *gper* open reading frame. Even if this transcript is translated, the resulting protein would lack large portions of transmembrane domains 1 and 2. The resulting protein would not fold properly and would not be properly integrated into the cell membrane and is likely to be completely devoid of function. We cannot exclude the possibility that genetic compensation is occurring to rescue phenotypes in the *gper^uab102^* deletion mutants [46]. In genetic compensation, upregulation of related genes occurs in response to a gene knockout. In the case of *gper*, there are no identified ohnologues or related genes in the zebrafish genome that are obvious candidates for genetic compensation. *gper^uab102^* embryos were shown to have a heart rate phenotype [2]. Thus, if genetic compensation is occurring, it is compensating for an adult phenotype but not an embryonic phenotype.

Our results suggest that GPER is not required for normal sex differentiation, gonad development or gonad function in zebrafish. These findings are consistent with studies from *Gper* mutant mice, which also reported no gonad development phenotypes [32, 33]. While GPER has been demonstrated to regulate the development and function of non-gonadal tissues [2, 47-49], we conclude that, in contrast to nuclear estrogen receptors, GPER is dispensable for gonad formation and function.

---

**Supplemental Table 1.**
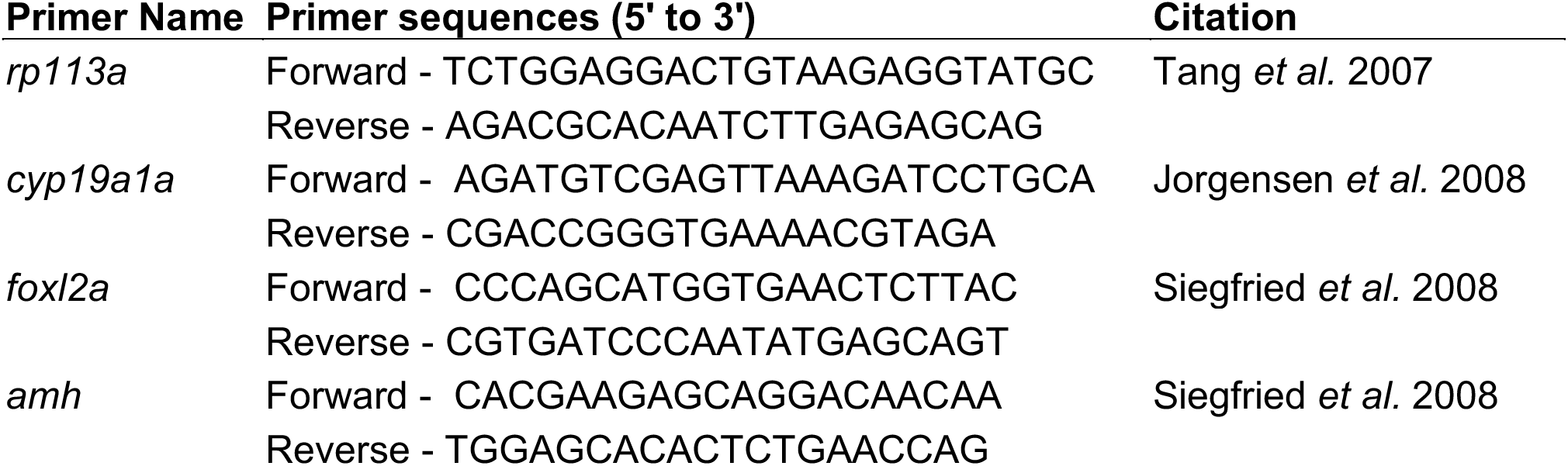
List of primers used for qRT-PCR.

## ACKNOWLEDGMENTS

We thank J. L. King for excellent technical assistance and S. C. Farmer and staff at the UAB Zebrafish Research Facility for animal care. This work was funded by the National Institutes of Health (National Institute of General Medical Sciences grants K12GM088010 to C.M.C., T32GM008111 to S.N.R., National Institute of Environmental Health Sciences grant R01ES026337 to D.A.G.).

## AUTHOR CONTRIBUTIONS

CMC and DAG conceived and designed the experiments, CMC performed the experiments, CMC and DAG analyzed/interpreted the data, SNR provided new tools/reagents, CMC and DAG wrote the manuscript. All authors approved the final manuscript.

## References

1. Lu H, Cui Y, Jiang L, Ge W. Functional Analysis of Nuclear Estrogen Receptors in Zebrafish Reproduction by Genome Editing Approach. Endocrinology 2017; 158:2292–2308.

2. Romano SN, Edwards HE, Souder JP, Ryan KJ, Cui X, Gorelick DA. G protein-coupled estrogen receptor regulates embryonic heart rate in zebrafish. PLoS Genet 2017; 13:e1007069.

3. Hoffman EJ, Turner KJ, Fernandez JM, Cifuentes D, Ghosh M, Ijaz S, Jain RA, Kubo F, Bill BR, Baier H, Granato M, Barresi MJF, et al. Estrogens Suppress a Behavioral Phenotype in Zebrafish Mutants of the Autism Risk Gene, CNTNAP2. Neuron 2016; 89:725–733.

4. Gorelick DA, Iwanowicz LR, Hung AL, Blazer VS, Halpern ME. Transgenic zebrafish reveal tissue-specific differences in estrogen signaling in response to environmental water samples. Environ Health Perspect 2014; 122:356–362.

5. Pang Y, Thomas P. Role of G protein-coupled estrogen receptor 1, GPER, in inhibition of oocyte maturation by endogenous estrogens in zebrafish. Dev Biol 2010; 342:194–206.

6. Carroll KJ, Esain V, Garnaas MK, Cortes M, Dovey MC, Nissim S, Frechette GM, Liu SY, Kwan W, Cutting CC, Harris JM, Gorelick DA, et al. Estrogen defines the dorsal-ventral limit of VEGF regulation to specify the location of the hemogenic endothelial niche. Dev Cell 2014; 29:437–453.

7. Bardet PL, Horard B, Robinson-Rechavi M, Laudet V, Vanacker JM. Characterization of oestrogen receptors in zebrafish (Danio rerio). J Mol Endocrinol 2002; 28:153–163.

8. Lassiter CS, Kelley B, Linney E. Genomic structure and embryonic expression of estrogen receptor beta a (ERbetaa) in zebrafish (Danio rerio). Gene 2002; 299:141–151.

9. Menuet A, Pellegrini E, Anglade I, Blaise O, Laudet V, Kah O, Pakdel F. Molecular characterization of three estrogen receptor forms in zebrafish: binding characteristics, transactivation properties, and tissue distributions. Biol Reprod 2002; 66:1881–1892.

10. Revankar CM, Cimino DF, Sklar LA, Arterburn JB, Prossnitz ER. A Transmembrane Intracellular Estrogen Receptor Mediates Rapid Cell Signaling. Science 2005; 307:1625–1630.

11. Thomas P, Pang Y, Filardo EJ, Dong J. Identity of an estrogen membrane receptor coupled to a G protein in human breast cancer cells. Endocrinology 2005; 146:624–632.

12. Liu X, Zhu P, Sham KW, Yuen JM, Xie C, Zhang Y, Liu Y, Li S, Huang X, Cheng CH, Lin H. Identification of a membrane estrogen receptor in zebrafish with homology to mammalian GPER and its high expression in early germ cells of the testis. Biol Reprod 2009; 80:1253–1261.

13. Ribas L, Valdivieso A, Diaz N, Piferrer F. Appropriate rearing density in domesticated zebrafish to avoid masculinization: links with the stress response. J Exp Biol 2017; 220:1056–1064.

14. Ribas L, Liew WC, Diaz N, Sreenivasan R, Orban L, Piferrer F. Heat-induced masculinization in domesticated zebrafish is family-specific and yields a set of different gonadal transcriptomes. Proc Natl Acad Sci U S A 2017; 114: e941–E950.

15. Ribas L, Valdivieso A, Diaz N, Piferrer F. Response to “The importance of controlling genetic variation - remarks on ‘Appropriate rearing density in domesticated zebrafish to avoid masculinization: links with the stress response’”. J Exp Biol 2017; 220:4079–4080.

16. Wilson CA, High SK, McCluskey BM, Amores A, Yan YL, Titus TA, Anderson JL, Batzel P, Carvan MJ, 3rd, Schartl M, Postlethwait JH. Wild sex in zebrafish: loss of the natural sex determinant in domesticated strains. Genetics 2014; 198:1291–1308.

17. Siegfried KR, Nusslein-Volhard C. Germ line control of female sex determination in zebrafish. Dev Biol 2008; 324:277–287.

18. Liew WC, Bartfai R, Lim Z, Sreenivasan R, Siegfried KR, Orban L. Polygenic sex determination system in zebrafish. PLoS One 2012; 7:e34397.

19. Chen W, Liu L, Ge W. Expression analysis of growth differentiation factor 9 (Gdf9/gdf9), anti-müllerian hormone (Amh/amh) and aromatase (Cyp19a1a/cyp19a1a) during gonadal differentiation of the zebrafish, Danio rerio. Biology of Reproduction 2017; 96:401–413.

20. Yang Y-J, Wang Y, Li Z, Zhou L, Gui J-F. Sequential, Divergent, and Cooperative Requirements of Foxl2a and Foxl2b in Ovary Development and Maintenance of Zebrafish. Genetics 2017; 205:1551–1572.

21. Rodriguez-Mari A, Yan YL, Bremiller RA, Wilson C, Canestro C, Postlethwait JH. Characterization and expression pattern of zebrafish Anti-Mullerian hormone (Amh) relative to sox9a, sox9b, and cyp19a1a, during gonad development. Gene Expression Patterns 2005; 5:655–667.

22. Dranow DB, Hu K, Bird AM, Lawry ST, Adams MT, Sanchez A, Amatruda JF, Draper BW. Bmp15 is an oocyte-produced signal required for maintenance of the adult female sexual phenotype in zebrafish. PLoS genetics 2016; 12:e1006323.

23. Schulz RW, Bogerd J, Male R, Ball J, Fenske M, Olsen LC, Tyler CR. Estrogen-induced alterations in amh and dmrt1 expression signal for disruption in male sexual development in the zebrafish. Environ Sci Technol 2007; 41:6305–6310.

24. Wang XG, Orban L. Anti-Mullerian hormone and 11 beta-hydroxylase show reciprocal expression to that of aromatase in the transforming gonad of zebrafish males. Dev Dyn 2007; 236:1329–1338.

25. Lin Q, Mei J, Li Z, Zhang X, Zhou L, Gui JF. Distinct and Cooperative Roles of amh and dmrt1 in Self-Renewal and Differentiation of Male Germ Cells in Zebrafish. Genetics 2017; 207:1007–1022.

26. Orn S, Yamani S, Norrgren L. Comparison of vitellogenin induction, sex ratio, and gonad morphology between zebrafish and Japanese medaka after exposure to 17alpha-ethinylestradiol and 17beta-trenbolone. Arch Environ Contam Toxicol 2006; 51:237–243.

27. Pang Y, Thomas P. Involvement of estradiol-17beta and its membrane receptor, G protein coupled receptor 30 (GPR30) in regulation of oocyte maturation in zebrafish, Danio rario. Gen Comp Endocrinol 2009; 161:58–61.

28. Peyton C, Thomas P. Involvement of epidermal growth factor receptor signaling in estrogen inhibition of oocyte maturation mediated through the G protein-coupled estrogen receptor (Gper) in zebrafish (Danio rerio). Biol Reprod 2011; 85:42–50.

29. Zhu Y, Liu D, Shaner ZC, Chen S, Hong W, Stellwag EJ. Nuclear progestin receptor (pgr) knockouts in zebrafish demonstrate role for pgr in ovulation but not in rapid non-genomic steroid mediated meiosis resumption. Front Endocrinol (Lausanne) 2015; 6:37.

30. Aizen J, Pang Y, Harris C, Converse A, Zhu Y, Aguirre MA, Thomas P. Roles of progesterone receptor membrane component 1 and membrane progestin receptor alpha in regulation of zebrafish oocyte maturation. Gen Comp Endocrinol 2018; 263:51–61.

31. Wang C, Prossnitz ER, Roy SK. G protein-coupled receptor 30 expression is required for estrogen stimulation of primordial follicle formation in the hamster ovary. Endocrinology 2008; 149:4452–4461.

32. Otto C, Fuchs I, Kauselmann G, Kern H, Zevnik B, Andreasen P, Schwarz G, Altmann H, Klewer M, Schoor M, Vonk R, Fritzemeier KH. GPR30 does not mediate estrogenic responses in reproductive organs in mice. Biol Reprod 2009; 80:34–41.

33. Isensee J, Meoli L, Zazzu V, Nabzdyk C, Witt H, Soewarto D, Effertz K, Fuchs H, Gailus-Durner V, Busch D, Adler T, de Angelis MH, et al. Expression pattern of G protein-coupled receptor 30 in LacZ reporter mice. Endocrinology 2009; 150:1722–1730.

34. Majumder S, Das S, Moulik SR, Mallick B, Pal P, Mukherjee D. G-protein coupled estrogen receptor (GPER) inhibits final oocyte maturation in common carp, Cyprinus carpio. Gen Comp Endocrinol 2015; 211:28–38.

35. Westerfield M. The Zebrafish Book. A Guide for the Laboratory Use of Zebrafish (Danio Rerio). Eugene, OR: University of Oregon Press; 2000.

36. Crowder CM, Lassiter CS, Gorelick DA. Nuclear Androgen Receptor Regulates Testes Organization and Oocyte Maturation in Zebrafish. Endocrinology 2018; 159:980–993.

37. Tang R, Dodd A, Lai D, McNabb WC, Love DR. Validation of zebrafish (Danio rerio) reference genes for quantitative real-time RT-PCR normalization. Acta biochimica et biophysica Sinica 2007; 39:384–390.

38. Jorgensen A, Andersen O, Bjerregaard P, Rasmussen LJ. Identification and characterisation of an androgen receptor from zebrafish Danio rerio. Comp Biochem Physiol C Toxicol Pharmacol 2007; 146:561–568.

39. Livak KJ, Schmittgen TD. Analysis of relative gene expression data using real-time quantitative PCR and the 2(-Delta Delta C(T)) Method. Methods 2001; 25:402–408.

40. Crowder CM, Lassiter CS, Gorelick DA. Nuclear androgen receptor regulates testes organization and oocyte maturation in zebrafish. Endocrinology 2017.

41. Parichy DM, Elizondo MR, Mills MG, Gordon TN, Engeszer RE. Normal table of postembryonic zebrafish development: staging by externally visible anatomy of the living fish. Dev Dyn 2009; 238:2975–3015.

42. Turola E, Petta S, Vanni E, Milosa F, Valenti L, Critelli R, Miele L, Maccio L, Calvaruso V, Fracanzani AL, Bianchini M, Raos N, et al. Ovarian senescence increases liver fibrosis in humans and zebrafish with steatosis. Dis Model Mech 2015; 8:1037–1046.

43. Fitzgerald AC, Peyton C, Dong J, Thomas P. Bisphenol A and Related Alkylphenols Exert Nongenomic Estrogenic Actions Through a G Protein-Coupled Estrogen Receptor 1 (Gper)/Epidermal Growth Factor Receptor (Egfr) Pathway to Inhibit Meiotic Maturation of Zebrafish Oocytes. Biol Reprod 2015.

44. Pang Y, Dong J, Thomas P. Estrogen signaling characteristics of Atlantic croaker G protein-coupled receptor 30 (GPR30) and evidence it is involved in maintenance of oocyte meiotic arrest. Endocrinology 2008; 149:3410–3426.

45. Pang Y, Thomas P. Role of natriuretic peptide receptor 2-mediated signaling in meiotic arrest of zebrafish oocytes and its estrogen regulation through G protein-coupled estrogen receptor (Gper). General and comparative endocrinology 2018.

46. Rossi A, Kontarakis Z, Gerri C, Nolte H, Holper S, Kruger M, Stainier DY. Genetic compensation induced by deleterious mutations but not gene knockdowns. Nature 2015; 524:230–233.

47. Natale CA, Duperret EK, Zhang J, Sadeghi R, Dahal A, O’Brien KT, Cookson R, Winkler JD, Ridky TW. Sex steroids regulate skin pigmentation through nonclassical membrane-bound receptors. Elife 2016; 5.

48. Meyer MR, Fredette NC, Daniel C, Sharma G, Amann K, Arterburn JB, Barton M, Prossnitz ER. Obligatory role for GPER in cardiovascular aging and disease. Sci Signal 2016; 9:ra105.

49. Ruiz-Palmero I, Hernando M, Garcia-Segura LM, Arevalo MA. G protein-coupled estrogen receptor is required for the neuritogenic mechanism of 17beta-estradiol in developing hippocampal neurons. Mol Cell Endocrinol 2013; 372:105–115.

